# Cryoablation temperature monitoring with dense ultrasonic speed-of-sound shift imaging

**DOI:** 10.64898/2026.01.04.697528

**Authors:** Gaya Lamm, Tal Grutman, Mike Bismuth, Tali Ilovitsh

## Abstract

Accurate temperature monitoring during cryoablation, a minimally invasive technique that destroys tissue locally by forming an ice ball around an inserted cryoprobe, is vital for achieving complete ablation while protecting surrounding tissue. We present a dense slowness-shift imaging method that estimates local speed-of-sound changes from ultrasound B-mode images using optical flow. This single-transducer, image-based approach enables mapping of spatial temperature change without requiring additional hardware. Cryoablation experiments in a tissue-mimicking phantom and ex vivo turkey breast demonstrated that slowness deviation increases with decreasing temperature. In the phantom, the dependence was linear (α_a_ = -20.70 ηs·m^−1^C°^−1^), while in turkey breast it presented an exponential relationship (α_t_ = 34.04×exp(0.075(-ΔT)) ηs·m^−1^C°^−1^). The algorithm detected sub-degree temperature variations and accurately tracked cooling down to -39.4 ± 5.6 °C. This work demonstrates the feasibility of ultrasound-based, noninvasive temperature monitoring during cryoablation, providing a scalable, real-time alternative to existing invasive or high-cost thermal assessment techniques.

## 1. Introduction

Cryoablation is a minimally invasive procedure used to destroy pathological tissues in a target area by exposing them to extremely low temperatures. This technique involves controlled freezing in a defined region, offering a less invasive alternative to surgical resection with reduced associated morbidity (Baust et al., 2007; Erinjeri & Clark, 2010; Gage & Baust, 1998; W. H. Yang et al., 2004). This targeted freezing process leads to irreversible cellular damage and tissue necrosis. Freezing injuries range from minor to severe resulting in different biological outcomes; minor injuries may trigger only an inflammatory response, whereas severe injuries involve the formation of intracellular and extracellular ice crystals. These ice crystals disrupt cellular membranes and organelles, ultimately resulting in apoptosis and necrosis. The extent of tissue injury depends on factors such as the cooling rate, the minimum temperature achieved, typically ranging from -20°C to -40°C, and the duration of exposure (Gage & Baust, 1998; W. H. Yang et al., 2004; Yiu et al., 2007).

Clinically, cryoablation is widely used to treat various conditions, including renal tumors, liver metastases, breast cancer, and cardiac arrhythmias (Baust et al., 2007; Bisbee et al., 2022; Bredikis & Wilber, 2012; Galati, Marra, et al., 2024; Khanmohammadi et al., 2023; Niu et al., 2014; Seager et al., 2020; M. Yang et al., 2025). As a minimally invasive technique, accurately defining and monitoring the treated region is critical for effective and safe treatment. During cryoablation, a thermal gradient is established around the cryoprobe, such that with increasing distance the achieved minimum temperature increases (Baust et al., 2007; Gage & Baust, 1998). This decrease in cooling efficiency has been demonstrated in previous studies (Hinshaw et al., 2010; Takami et al., 2015). This spatial variation in temperature influences the shape of the frozen region (encapsulated by the freeze margin) and the surrounding hyperemic tissue, both of which must be considered to ensure complete destruction of pathological cells while preserving healthy structures (Baust et al., 2007; Gage & Baust, 1998; Galati, Marra, et al., 2024; Thaokar & Rabin, 2012). Therefore, temperature monitoring is essential to confirm that a lethal temperature threshold is reached within the target area, and to avoid excessive freezing of adjacent tissue.

Currently, a common method for temperature monitoring during cryoablation includes the use of thermocouples. In research settings, invasive temperature sensors such as fiber optic sensors, thermistors, and resistance temperature detectors have been used for monitoring temperature and freezing depth (Ikiades et al., 2023; Martin et al., 2017; Roriz et al., 2020; Schena et al., 2016; Spasopoulos et al., 2020), though their clinical adaptation remains limited. These sensors are inherently invasive and provide only localized, point-based temperature measurements rather than a full spatial temperature map of the treated region, which restricts their ability to capture thermal gradients or heterogeneity across the treated region. Infrared (IR) thermometry has also been explored in research as a non-contact method to estimate surface temperatures during cryoablation (Pogrel et al., 1996; Yan et al., 2007). However, IR thermometry is limited to superficial measurements and cannot capture temperature dynamics within deeper tissue, since IR radiation does not penetrate beneath the surface. To overcome this limitation, magnetic resonance imaging thermometry has been employed as a non-invasive method for visualizing temperature changes throughout the tissue volume. MRI-guided cryoablation has been investigated in various studies (Bomers et al., 2013; de Jager et al., 2023; De Marini et al., 2021; Gangi et al., 2012; X. Liu et al., 2012; Morrison et al., 2008; Overduin et al., 2016; Sidana et al., 2024; Tuncali et al., 2007) and is increasingly used in clinical settings due to its ability to provide spatially resolved temperature information. Despite its advantage as a reliable tool for visualizing the ablated tissue and its surrounding region without invasive contact, MRI-guided monitoring introduces a high cost, and requires specialized infrastructure, thus limiting availability.

Ultrasound (US) is an especially appealing technology for temperature monitoring, as it is noninvasive, cost effective and capable of imaging deep tissues in real time (Abiteboul & Ilovitsh, 2022; Karlinsky et al., 2024). US imaging operates by transmitting acoustic waves into tissue and detecting the echoes that return from interfaces with different acoustic impedances, allowing the reconstruction of structural information based on the amplitude and timing of these reflections. It is widely utilized for guiding and monitoring thermal ablation procedures due to its real-time feedback, portability, and wide availability (de Jager et al., 2023; De Marini et al., 2021; Ding et al., 2016; Foiret & Ferrara, 2015; Grutman & Ilovitsh, 2023; D. Liu & Ebbini, 2010; Rohfritsch et al., 2023; Sehgal et al., 1986; Seip & Ebbini, 1995; Sidana et al., 2024; Simon et al., 1998; Zhou et al., 2019). US is also used for guiding and monitoring cryoablation procedures, although its role is still evolving. During cryoablation, the formation of ice leads to a hyperechoic region with posterior acoustic shadowing, which can obscure the visualization of deeper tissue structures. Despite this limitation, the appearance of the hyperechoic zone serves as an indirect indicator of the frozen region (Galati, Pasculli, et al., 2024; Gilbert et al., 1985; Laugier & Berger, 1993; Onik et al., 1984; Pfleiderer et al., 2002, 2005; Poplack et al., 2015; Tacke et al., 1999; Thai et al., 2023; Ward et al., 2019). However, this information is qualitative rather than quantitative; US imaging cannot reveal the actual temperature within the ice ball or in the surrounding tissue, and lethal isotherms such as -20°C or -40°C cannot be distinguished from visual appearance alone. As a result, clinicians performing cryoablation must rely solely on the B-mode image without access to spatially resolved thermal information, limiting their ability to assess treatment adequacy or avoid unintended cooling of adjacent structures. To enhance the assessment of thermal and structural changes during cryoablation, researchers have explored advanced ultrasound techniques. Ultrasound elastography, for instance, measures tissue stiffness and has been investigated for monitoring the frozen zone post-ablation (Dall’Alba et al., 2012).

Another promising approach involves estimating the local speed-of-sound (SoS), which is sensitive to temperature-dependent changes in tissue properties. SoS measurements were utilized in various applications such as estimating ice-content, assessing thaw cycles, and identifying pre-thaw stages in frozen foods (Aparicio et al., 2008; Chang et al., 2023; Gallo et al., 2018; Jha et al., 2023) as well as temperature estimation in orange juice (Carcione et al., 2007). A comprehensive review by Liang et al., (2024) examined acoustic properties measured in materials down to -50°C, including water, aqueous solutions, lipids and biological tissues. A pronounced change in SoS was observed (i.e., longitudinal velocity) in water, aqueous solutions and biological tissues when the samples underwent freezing, whereas lipids and oils exhibited a more gradual but still distinct shift. Contrary to the well-studied linear relationship of US image strain and temperature change in heating applications of biological tissue (Civale et al., 2013; Grutman & Ilovitsh, 2023; Miller et al., 2002), it has been shown that tissue composition significantly affects SoS behavior during freezing, particularly the water-to-lipid ratio in biological tissues (Liang et al., 2024). In water-rich samples, the SoS change is dominated by the phase transition of water, while in biological tissues, additional factors such as fat distribution and microstructural freezing dynamics cause smaller SoS changes (Liang et al., 2024). Consequently, it is expected that the relationship will be more complex in ex-vivo tissues containing lipids, compared to water-dominated samples. Studies reviewed by (Liang et al., 2024) relied on pulse-echo or through-transmission configurations combined with sample-insertion or multiple-reflection methods, all of which estimate SoS through time-of-flight measurements. Importantly, these approaches require two transducers or a reflector, provide single-point or bulk measurements, and are generally performed in controlled laboratory setups, with freezing induced by air-cooling or plate-based conduction rather than clinically relevant cryoprobes. As a result, prior SoS-based methods are not directly applicable to spatially resolved, real-time temperature monitoring during in-situ cryoablation.

These findings establish a foundation for employing SoS shifts as indicators of underlying acoustic property changes in tissue during cryoablation. In particular, the strong dependence of SoS on temperature enables the estimation of local temperature variations based on measured shifts in sound speed. Here, we propose a dense SoS-shift imaging (DSI) algorithm for SoS shift estimation, which was implemented in an experimental platform designed to enable controlled cryoablation in different samples. Simultaneous imaging and recording of ground-truth temperature was conducted with a thermocouple monitoring temperature during acquisitions of raw RF data. Our algorithm transforms the raw data into B-mode US images and analyzes them for pixel displacements induced by temperature variations between 20°C and -40°C, using the Farnebäck optical flow algorithm (Farnebäck, 2003). A regularized inverse problem framework was subsequently established to relate these displacements to SoS shifts. These shifts correspond to temperature decreases and can be utilized for estimating temperature (Figure 1). This algorithm was originally developed by our group for monitoring thermal ablation and is here adapted for cryoablation (Grutman & Ilovitsh, 2023). This framework allowed us to directly characterize the physical relationship between cryogenic temperature decrease and slowness deviation of SoS in both tissue-mimicking phantoms and ex vivo tissue samples. By establishing this quantitative SoS-temperature dependence, including material-specific behaviors and their functional forms, we demonstrate how image-based slowness estimation can be used to infer temperature profiles during cryoablation. This non-invasive approach offers a potential alternative to existing monitoring techniques, enabling deep-tissue assessment using conventional ultrasound systems and reducing reliance on implantable sensors or costly imaging infrastructure.

**Figure 1.**
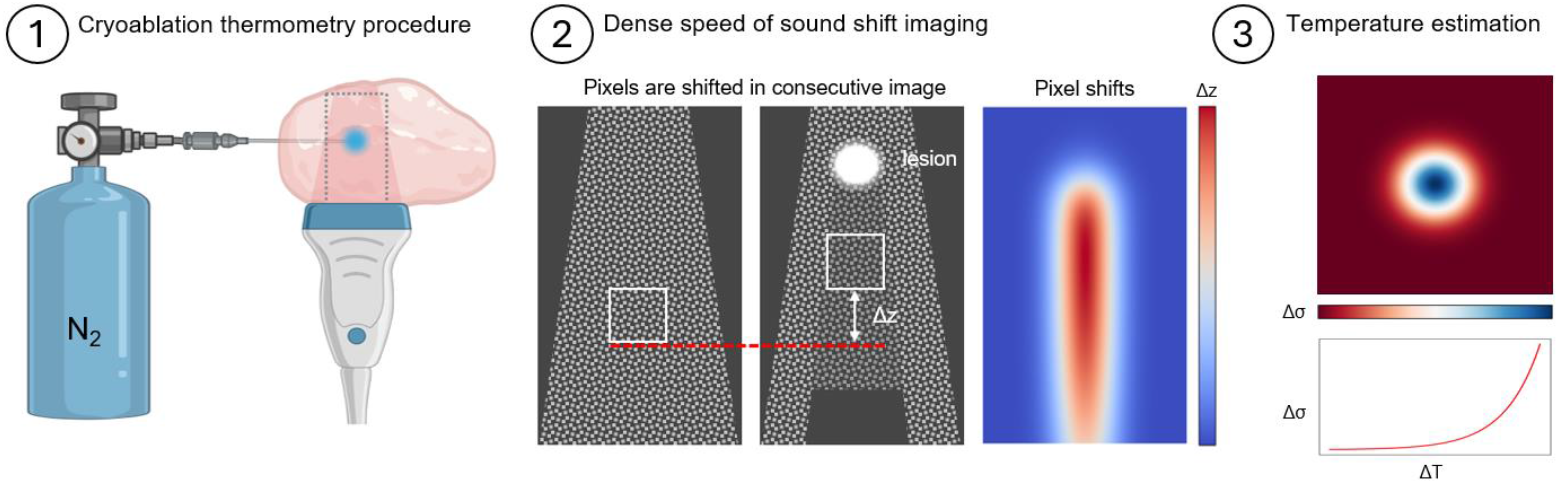
Cryoablation thermometry illustration. (1) Cryoablation is performed on a sample using a liquid nitrogen tank connected to a cryoprobe, which injects liquid nitrogen to locally cool the sample. An US imaging transducer is positioned perpendicular to the sample and records RF data during the cryoablation procedure. (2) RF data are converted into B-mode images and pixel displacements within the region affected by the growing cryoablation lesion are tracked using optical flow. (3) Pixel displacements between consecutive frames are used to compute the local slowness deviation within the imaged region. A calibration between slowness and temperature is established using simultaneous thermocouple measurements.

## 2. Theory

The mathematical relationship between the predicted echo-shift or change in round-trip time δτ, and the temperature change δT along the wave propagation path, was described in our previous work (Grutman & Ilovitsh, 2023). Briefly, the relationship between of δτ to the local shift in the sound speed is given by:

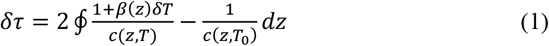

where the difference in round-trip time δτ is the cumulative sum of the inverse of changes in local speed of sound c(z,T) due to temperature change δT from the initial temperature T_0_ in tissue with a thermal expansion coefficient of β(z) along the propagation path. In our method, the approach is linearization of the problem, such that equation (1) can be vectorized, and the following equation is obtained:

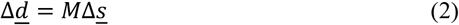

where the pixel shift Δ<u>d</u> is observed in a plane-wave B-mode image due to an echo-shift of δτ. Δ<u>s</u> represents the change in sound slowness, described by:

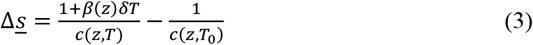

Once the integral operand Δ<u>s</u> is isolated, it is equivalent to the derivative of (1) such that

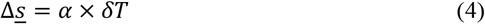

where α is a medium-specific parameter describing the relationship of speed of sound with temperature, as well as a unit conversion parameter from strain to temperature. Importantly, in this study α may be a function of temperature change, ΔT, rather than a constant coefficient, such that the relationship between Δ<u>s</u> and ΔT is described by α(ΔT) and may not be linear. This is due to the varying acoustic properties in low temperatures that are dependent on tissue composition, as described by (Liang et al., 2024). α is extracted from a curve of Δs, which is calculated by the algorithm, vs. ΔT, which is measured as ground-truth. This calibration is common to thermal measurements from acoustic strain images (Chen et al., 2025; Foiret & Ferrara, 2015; Obara et al., 2021; Sheng et al., 2025).

To calculate the change in sound slowness Δs, the following inverse problem is formulated:

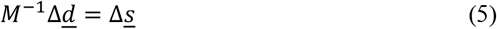

The pixel shift Δ<u>d</u> is measured from acquired B-mode images. The optimal Δs which satisfies the data is calculated via a Tikhonov pseudo-inverse. The spatial gradient was regularized using L2 regularization of the local gradient. Finally, the change in sound slowness Δ<u>s</u> is estimated by solving the following least squares problem, using M^†^ as a pseudo-inverse matrix for the solution (Grutman & Ilovitsh, 2023; Stähli et al., 2020):

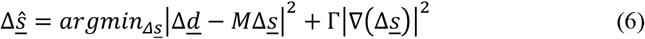

Ultrasound plane wave B-mode images are acquired, and the pixel shift in between consecutive images is computed via optical flow to produce the measured vector Δ<u>d</u>. In our previous work (Grutman & Ilovitsh, 2023), the optical-flow-based slowness deviation reconstruction method was developed for heating processes, where temperature increases lead to an increase in speed of sound. Thus, the direction of apparent speckle motion in B-mode images corresponded to positive speed of sound shifts. In contrast, during cryoablation, tissue cooling results in a decrease in SoS, which causes the speckle pattern to shift in the opposite direction. To maintain consistency with the inverse model and ensure correct speed of sound shift estimation, the sign of the computed pixel displacements field was inverted prior to solving the reconstruction problem. Then, the measurements vector Δ<u>d</u> was inverted as an inverse problem to receive the estimator for change in slowness, Δ<u>ŝ</u>. Finally, the change in slowness can be utilized for temperature estimation.

## 3. Methods

### 4.1 Cryoablation procedure and setup

Two types of samples were used. The first was a tissue mimicking phantom. The phantom was made by heating 4.5 grams of agarose powder (214010, Becton, Dickinson and Company, MD, USA) in 300 ml deionized water in the microwave in pulses until boiling. The mixture was left in room temperature for 10 minutes and stirred with a magnetic stirrer for another 10 minutes. 1.5 grams of silicone carbide (357391, Sigma Aldrich, MO, USA) were added to the mixture as acoustic scatterers and stirred for 30 minutes. The mixture was poured into a mold to congeal. The result was a phantom with a shallow sunken circular area as a target for nitrogen contact. The nitrogen was gently sprayed on the circular target area from above, while the imaging transducer, positioned perpendicularly and 4-5 mm under the sunken area plane, imaged the area adjacent to the sprayed target area. This area’s temperature decreased due to the ice-ball that was formed around the target area, as presented in Figure 2(c). The procedure was repeated in six samples of this type.

**Figure 2.**
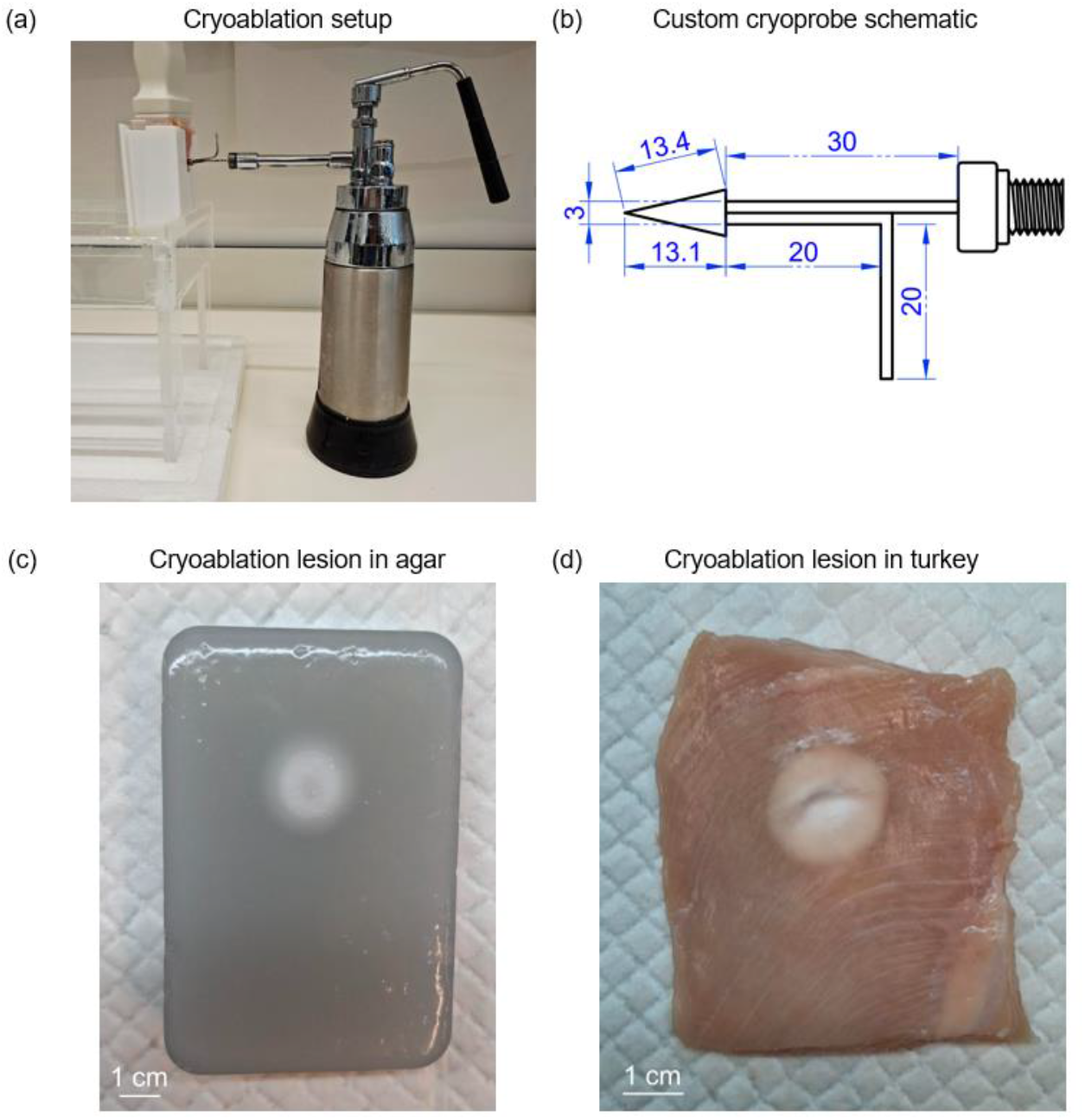
Experimental setup and samples used for cryoablation experiments. (a) Experimental setup, showing the US imaging transducer positioned perpendicular to the cryoprobe, cryoprobe connected to a liquid nitrogen tank, and 3D printed white box as a sample holder, during cryoablation. (b) Schematic illustration of the custom-designed cryoprobe, consisting of a sharp copper tip connected to a stainless-steel pipe, enabling controlled surface cooling without direct nitrogen injection into the sample. Units are mm. (c) Tissue-mimicking agar phantom after local cryoablation, exhibiting the frozen area formed at the target surface exposed to the cryoprobe cooling. (d) Ex vivo turkey breast sample following local cryoablation using the cryoprobe, showing the frozen region at the contact site.

The second set of samples were fresh ex-vivo turkey breast samples, obtained from a local butcher shop. Samples were cut from the thickest part of a turkey breast in a single piece that could fill the field of view and contain the cryoprobe depth. Each sample was placed in a 3D printed cartridge, and the custom cryoprobe was inserted into it to a depth of 4-5 mm. To minimize reflections, 6 mm rubber was placed around the samples and coupled with ultrasound gel. The experimental setup is presented in Figure 2(a). As a ground truth, in both phantoms and turkey breasts, a T-type thermocouple connected to a thermometer (TL253, Kamtop, China) was placed within the frozen region, as close as possible to the cryoprobe. The temperature measured with the thermocouple was recorded at the same time points of each data acquisition. The procedure was repeated in five samples of this type.

Direct injection of liquid nitrogen into the sample can lead to rapid pressure buildup and consequential movements of the tissue. This is undesirable for reproducible thermal measurements based on US imaging. Therefore, it is required that the freezing system confine the liquid nitrogen in the probe and would not allow it to be released into the tissue. We designed a custom liquid nitrogen cryoprobe connected to a 350 ml stainless steel liquid nitrogen sprayer can (Mini cryo-can liquid nitrogen sprayer, Rana Traders, India) (Figure 2(a)). A custom freeze-head was fabricated to interface with the tank’s valve, allowing for the controlled release of nitrogen (Figure 2(b)). As the nitrogen expanded and vaporized at the temperature probe outlet, it absorbed heat from the probe structure via conduction, thereby reducing the probe’s temperature to cryogenic levels (Erinjeri & Clark, 2010). This cooling mechanism enabled effective and localized tissue freezing at the contact site, minimizing sample motion and imaging artifacts during ultrasound acquisition.

### 4.2 Ultrasound imaging

Imaging of the sample was performed using an L12-5 50 mm linear transducer (ATL Philips, WA, USA) positioned perpendicular to the cryoprobe and coupled with ultrasound gel. A programmable ultrasound system (Vantage 256, Verasonics Inc., WA, USA) was used for imaging of a 60 mm × 40 mm field of view. A plane steering acquisition protocol was used to acquire nine plane waves per image. Three main steering angles were selected at 5° intervals [-5°, 0°, 5°], each composed of three steering angles selected at 1.5° intervals (for example, [3.5°, 5°, 6.5°]). Raw RF data was acquired automatically every 9 seconds, for a duration of 2 minutes, while the sample underwent local cryoablation. The data was stored for offline processing.

### 4.3 Post processing and speed of sound shift calculation

RF data was beamformed and interpolated onto a 600 × 390 pixel^2^ grid, such that the axial z direction represents 60 mm and the lateral x direction represents 40 mm. Each image was then Hilbert transformed, and the 1.5° angled plane waves sub-sets were compounded such that each image acquisition provided three main steering angles. The dense optic flow was calculated between consecutive images at each of the main steering angles using Farneback’s method (Farnebäck, 2003) and was inverted to maintain consistency with the inverse model. The three calculated pixel-shift images acquired at each time interval were down-sampled ×5 in the lateral direction and ×10 in the axial direction, then stacked to create the data vector. The data vector Δ<u>d</u> was multiplied by the matrix M^†^, and the regularization parameters affecting smoothness were selected as Γ_x_=10 and Γ_z_ = Γ_xz_ = 1. Then, the image was up-sampled back to the original image dimensions and summed. To obtain a curve for slowness deviation vs. temperature change, six agar-phantom samples and five turkey-breast samples were used, one time each, in a cryoablation experiment. The data points for each sample were grouped by their temperature change (ΔT) value proximity, and the mean and standard deviation were calculated for each data point, both for ΔT and for their corresponding slowness deviation (Δσ) mean value. DSI was implemented in python, and the pixel displacements were calculated via the OpenCV library (Bradski, 2000). The inverse problem solution (Eq. 5) was implemented as a sparse matrix multiplication using scipy (Virtanen et al., 2020).

## 4. Results

### 5.1 Tissue mimicking phantom results

Initial experiments were conducted in a tissue-mimicking phantom. A thermocouple was inserted prior into the phantom in proximity to the center of the target area in the sample to a depth of 1 cm. The phantom underwent local cryoablation in a target area (methods), and an ice-ball was created around the contact site. In each repetition, the duration of the freezing treatment ranged from 1.5 to 2 minutes until the temperature decreased by over 50°C from the initial temperature (11.30 ± 1.13°C). Consecutive US images were acquired automatically every 9 seconds and the temperature measured by the thermocouple was simultaneously recorded. The temperature change is the subtraction of the temperature recorded at the time point from the initial temperature. Cooling was performed until a large ice-ball was created and the hyperechoic ice-ball edge was visible in the B-mode frame (Figure 3(a)), meaning the ice-ball grew and entered the transducer’s imaging plane, resulting in a dark region below the hyperechoic edge. Pixel shifts between consecutive US B-mode images were calculated (Figure 3(b)) and used to generate the map of slowness deviation at each point of the imaged region (Figure 3(c)).

**Figure 3.**
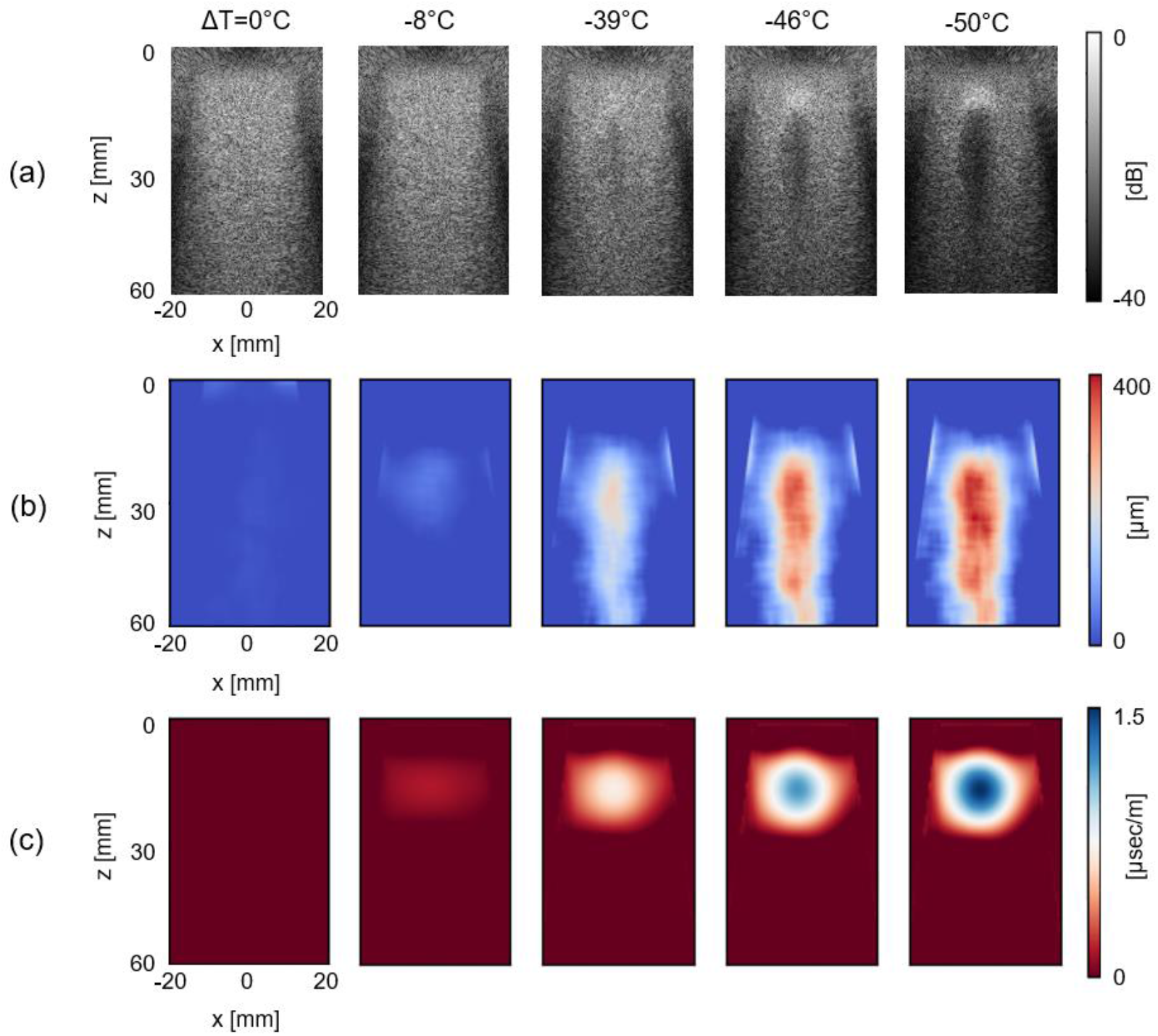
Effect of temperature decrease on the slowness deviation calculated by DSI in a tissue-mimicking sample. Cryoablation treatment was performed, and the temperature change was measured at each timepoint ΔT = 0°C, -8°C, -39°C, -46°C, -50°C. (a) B-mode images of the treated region at each timepoint. (b) Pixel shifts in the respective B-mode images, calculated by the algorithm. (c) Sound slowness calculated by the algorithm from the pixel shifts in (b). Axes are shown only for the first timepoint in (a)-(c) and are common to all timepoints. Colorbar is common to all subfigures in (a)-(c).

Slowness deviation was assessed as a function of temperature reduction (Figure 4(a)). The maximum slowness deviation measured within each imaged region was used to construct the slowness deviation-temperature curve. Since the largest temperature decrease was expected to occur in the area nearest to the cryoprobe and thermocouple, this region was assumed to exhibit the highest slowness deviation. A linear fit was then applied to the collected data using linear regression yielding R^2^ = 0.988, which describes the relationship between temperature change and slowness deviation with a slope of α_a_ = -20.70 ηs/mC° in agar phantom. In addition, the areas in the image where the temperature was below 0°C based on the calibration, were calculated in each image and are presented as a function of temperature (Figure 4(b)), starting from 121.0 ± 60.0 mm^2^ and gradually reaching 312.0 ± 85.4 mm^2^ as the recorded temperature decreased. This enables the estimation of the frozen region around the cryoprobe, complementary to the maximal temperature change estimation.

**Figure 4.**
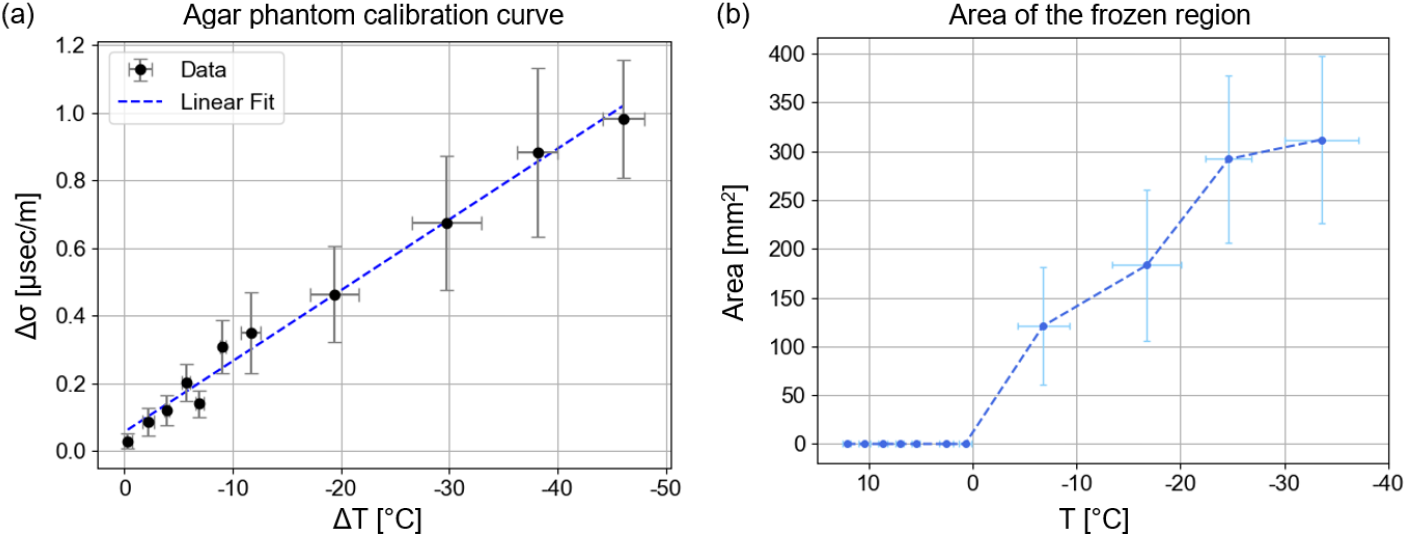
Analysis of temperature-slowness relationship and cold-region estimation during cryoablation in agar. (a) Calibration curve relating temperature change (ΔT) to slowness deviation (Δσ) during cryoablation in agar phantoms. The maximum slowness deviation measured in each imaged region was plotted against the corresponding temperature change recorded by the thermocouple. Data are presented as mean ± standard deviation in both axes (N = 6). A linear fit was applied using linear regression, yielding R^2^ = 0.988. (b) Area in reconstructed slowness deviation image where T<0°C as function of the temperature T recorded by the thermocouple. Data are presented as mean ± standard deviation in both axes (N = 6).

### 5.2 Ex-vivo turkey breast sample

A similar procedure was performed on an ex vivo turkey breast sample to validate the method in biological tissue. In this case, the freezing was achieved using the custom-made liquid nitrogen cryoprobe described earlier, which was inserted approximately 1 cm into the tissue. The thermocouple was positioned near the cryoprobe tip within the freezing zone to record local temperature changes. Cryoablation was applied for 1.5-2 minutes, producing a temperature drop of over 50°C from the initial temperature (15.30 ± 3.60°C) and forming a visible ice ball around the probe. US B-mode images (Figure 5(a)) were continuously acquired at 9-second intervals throughout the procedure, while the temperature was simultaneously recorded via the thermocouple. The resulting pixel shifts between consecutive B-mode frames were calculated (Figure 5(b)) and converted into slowness deviation maps (Figure 5(c)), following the same analysis pipeline used for the phantom experiments.

**Figure 5.**
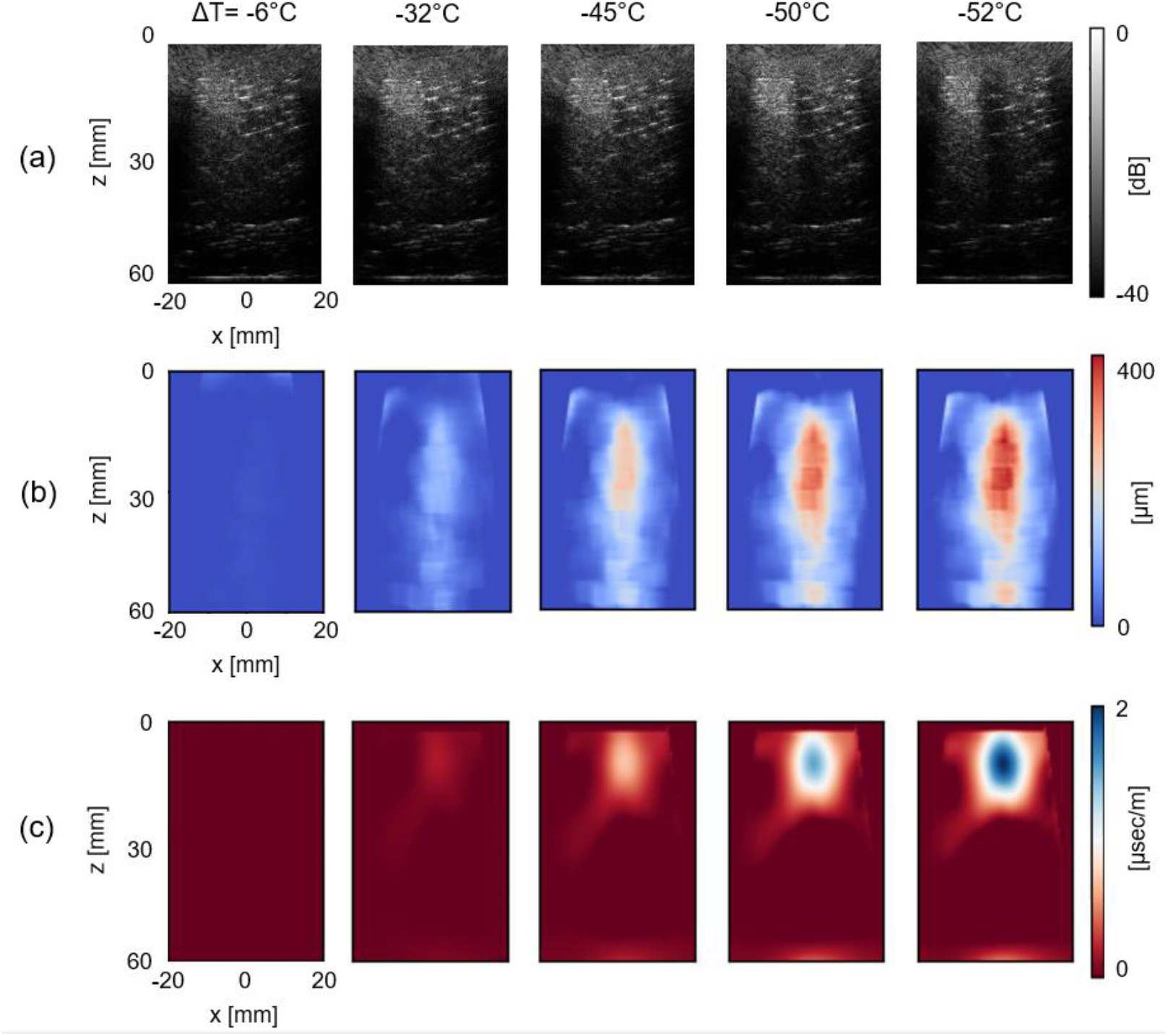
Effect of temperature decrease on the slowness deviation calculated by DSI in an ax-vivo turkey breast sample. Cryoablation treatment was performed, and the temperature change was measured at each timepoint ΔT = -6°C, -32°C, -45°C, -50°C, -52°C. (a) B-mode images of the treated region at each timepoint. (b) Pixel shifts in the respective B-mode images, calculated by the algorithm. (c) Sound slowness calculated by the algorithm from the pixel shifts in (b). Axes are shown only for the first timepoint in (a)-(c) and are common to all timepoints. Colorbar is common to all subfigures in (a)-(c).

The resulting data were analyzed following the same procedure as in the phantom experiment. The maximum slowness deviation from each region was correlated with temperature change. Since the data displayed an exponential relationship between slowness deviation and temperature change, the data was logarithmically transformed to obtain a linear relationship. Then, a linear fit was calculated using linear regression, yielding ln(Δσ)=0.075×(-ΔT)-3.38, R^2^ = 0.950. The resulting coefficients of the linear fit of the transformed data were taken as the coefficients of the exponential fit of the data, such that it is described by Δσ=34.04×exp(0.075×(-ΔT)) ηs/mC° (Figure 6(a)). The area where T<0°C was also calculated and is presented as a function of temperature (Figure 6(b)), showing an estimation of the frozen region, growing from 67.10 ± 13.90 mm^2^ and reaching 504.90 ± 127.20 mm^2^ at the end of the cryoablation process.

**Figure 6.**
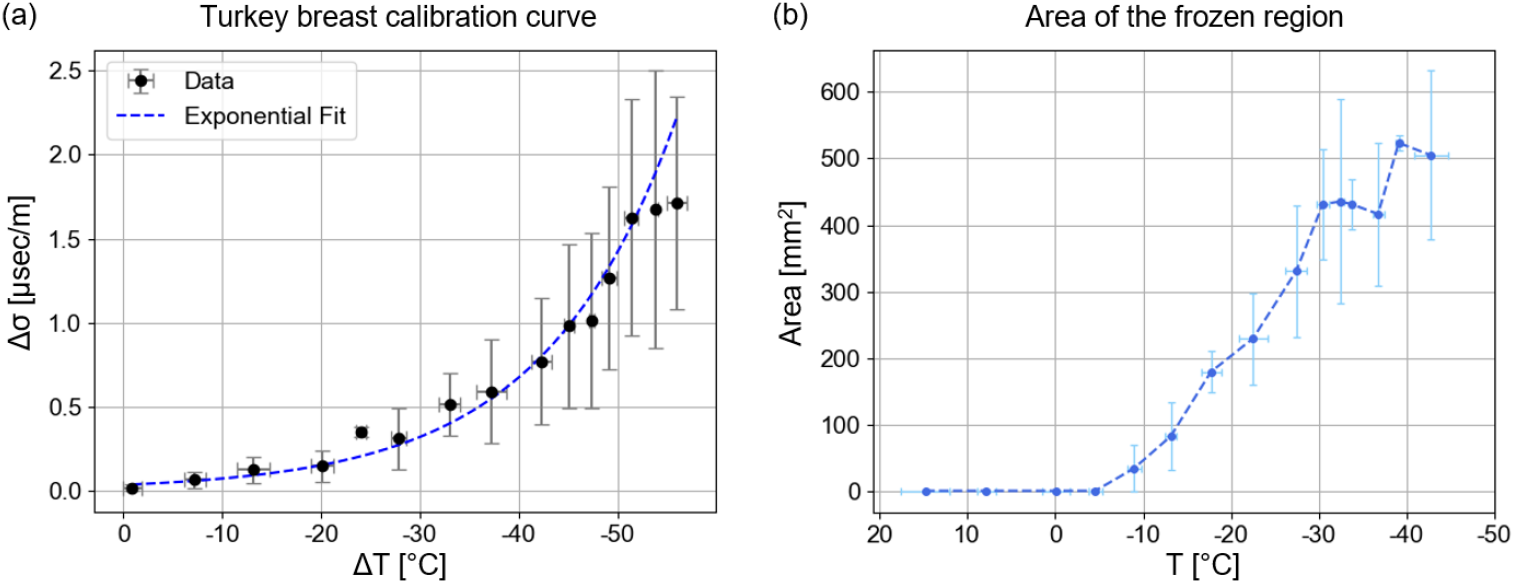
Analysis of temperature-slowness relationship and cold-region estimation during cryoablation in turkey breast. (a) Calibration curve relating temperature change (ΔT) to slowness deviation (Δσ) during cryoablation in turkey breast. The maximum slowness deviation measured in each imaged region was plotted against the corresponding temperature change recorded by the thermocouple. Data are presented as mean ± standard deviation in both axes (N = 5). (b) Area in reconstructed slowness deviation image where T<0°C as function of the temperature T recorded by the thermocouple. Data are presented as mean ± standard deviation in both axes (N = 5).

## 5. Discussion and Conclusions

In this study, we investigated the feasibility of using SoS shifts to monitor temperature distributions during cryoablation, motivated by the known sensitivity of SoS to temperature-dependent changes in biological tissues. Conventional SoS measurement approaches typically rely on two transducers or a transducer-reflector configuration and provide SoS or time-of-flight measurements on a single-point basis (Liang et al., 2024). Our method enables spatially resolved estimation of SoS shifts across the entire imaging region using a single transducer. This allows for real-time visualization of the cooling process and temperature gradients during cryoablation. Compared with thermal strain imaging, which estimates SoS changes indirectly through echo time shifts and often relies on cross-correlation between pre- and post-heating frames, our previously developed algorithm, DSI, (Grutman & Ilovitsh, 2023), directly computes the slowness deviation field from pixel displacements using optical flow. Previously we showed that DSI offers several key advantages over thermal strain imaging, including improved spatial coherence, greater robustness across both simulated and ex vivo data, and significantly faster computation by eliminating the correlation search step. These improvements make our algorithm well suited for real-time monitoring applications. Combined with a simple, low-cost cryogenic platform and an US imaging system, this approach provides a practical and scalable framework for spatially resolved temperature mapping during cryoablation.

Our method was implemented during cryoablation and calculated the slowness-deviation caused by temperature changes in the sample. We measured slowness deviations both in a tissue-mimicking phantom and in an ex vivo sample, at depths extending several millimeters below the surface adjacent to the cryoprobe contact point. These measurements revealed a consistent pattern of increased slowness deviation with decreasing temperature. While in tissue-mimicking agar phantom, a water-rich sample, the data exhibited a linear relationship between temperature change and slowness deviation, with a coefficient of α_a_ = -20.70 ηs/mC° (Figure 4(a)), in turkey breast sample, a biological tissue containing fat and other micro-structures, the relationship was exponential, described by α_t_ = 34.04×exp(0.075×(-ΔT)) ηs/mC° (Figure 6(a)). This indicates that the acoustic response to cooling, represented here by the α parameter, is material-specific and depends on both composition and structural properties such as water and lipid content, as expected (Liang et al., 2024). Therefore, α should be experimentally determined for each tissue type to ensure accurate temperature estimation during cryoablation.

Complementary to the temperature change estimation, it is also possible to estimate the frozen region around the cryoprobe using DSI. In agar phantom, the frozen region where the temperature was lower than 0°C was initially ×1.8 larger than in turkey breast, but the final size of the frozen region was ×1.62 larger in turkey breast compared to the agar phantom (Figure 4(b), Figure 6(b)). This difference could be the result of several parameters: the maximal temperature decrease achieved in each sample type (decrease of 46.1 ± 2.0°C from initial temperature 11.30 ± 1.13°C in agar vs. decrease of 56.10 ± 1.0°C from initial temperature 15.30 ± 3.60°C in turkey breast), which directly affects the temperature distribution in the surrounding tissue during the freezing process, and resulting in a larger frozen surrounding area for a greater decrease in temperature; the size of the contact area with the cryogenic temperature origin, resulting in a larger frozen area when the contact site size is larger, reached several mm in turkey breast, while in agar phantom ranged from 0.5-1 cm;, and the sample composition, which is more aqueous in agar, while richer in lipids and microstructures in turkey breast, affecting the rate of the ice ball growth and the temperature distribution (Yoon et al., 1998).

Although the ice ball itself appears as a hypoechoic region in B-mode ultrasound and attenuates the acoustic signal, preventing direct imaging beyond its boundary, our approach capitalizes on the measurable region adjacent to the ice ball. The B-mode image does not capture the pixel shifts beyond the hyperechoic boundary due to the strong attenuation and scattering at the frozen interface. However, by calibrating the slowness deviation in the adjacent region to temperature measurements, we were able to estimate the temperature profile indirectly inside the ice ball. This concept aligns with prior studies that profiled and modeled isotherms around cryoprobes, consistently showing that critical lethal isotherms such as -20°C and -40°C reside within the frozen volume (Hinshaw et al., 2010; Littrup et al., 2009; Shah et al., 2016). Thus, estimating the temperature in the surrounding tissue offers a noninvasive pathway for inferring thermal conditions within the ice ball itself.

Notably, frozen water expands, and this physical expansion may locally displace surrounding tissue as the ice ball grows. This mechanical deformation could contribute to pixel shifts observed in the B-mode image, in addition to those caused by temperature-dependent acoustic property changes. While such displacement does not affect traditional time-of-flight-based SoS measurements, it may influence our slowness-based approach, introducing a secondary effect that should be accounted for in future refinements of the method. Nevertheless, the observed slowness deviations remained consistent with the expected acoustic response to cooling, suggesting that the dominant contributor is indeed temperature change.

A key strength of our approach is its compatibility with standard US equipment and the use of a single imaging transducer. Although our algorithm was designed for real-time implementation, as it does not require complex calculations and has a runtime of 3 frames per second (Grutman & Ilovitsh, 2023), the current experiments were performed ex vivo and processed offline. Future work should therefore focus on real-time integration and in vivo validation, where physiological motion, perfusion, and acoustic heterogeneity may influence accuracy. Since DSI operates directly on beamformed B-mode images, it can be readily implemented with a wide range of transducers, including linear, curvilinear, and phased-array probes, by accounting for the image geometry and lateral sampling. Plane-wave acquisitions are particularly suitable, as they provide the high frame rates needed for robust optical-flow estimation.

The sensitivity of the algorithm depends primarily on the temporal spacing between consecutive B-mode frames. Because the method uses optical flow to estimate the pixel-wise motion between successive ultrasound images, shorter frame intervals improve sensitivity by capturing smaller and smoother displacements. This enhances the precision of the optical flow calculation and, consequently, the accuracy of the derived slowness deviation. In the present implementation, using data acquired at 9-second intervals, DSI detected slowness deviations corresponding to temperature variations ranging from sub-degree changes to several degrees. In the context of cryoablation, where tissue temperatures typically decrease from physiological levels (∼37°C) down to -40°C or lower, such sensitivity is sufficient to monitor both the progression of freezing in the target zone and to ensure that surrounding tissues are not inadvertently overcooled. In this study, the lowest temperature achieved ex vivo was -39.4 ± 5.6°C, and the corresponding slowness deviation was successfully detected by the algorithm. The observed temperature decrease was strongly correlated with both the cooling duration and the growth of the ice ball surrounding the probe. However, as the ice ball enlarges, it creates a hyperechoic region that limits US penetration, imposing a practical constraint on the minimal temperature that can be measured. The formation and evolution of the ice ball during cryoablation depend on the probe’s cooling rate and the thermal properties of the surrounding tissue (Littrup et al., 2009; W. H. Yang et al., 2004). A commercial cryoprobe, capable of reaching cryogenic temperatures within a shorter period, would likely reduce excessive ice-ball formation and enable detection of even lower temperatures when combined with a shorter frame acquisition interval.

In summary, our algorithm enables spatially resolved, noninvasive monitoring of temperature during cryoablation, capturing slowness deviations across the imaging region and providing indirect estimates of frozen tissue temperatures. The method performed robustly in both phantoms and ex vivo samples and can be adapted for real-time use with various ultrasound transducers. Calibration of the α parameter is needed for different tissues when implementing this method for cryoablation procedures. Future work should extend this approach to in vivo settings and optimize acquisition parameters to enhance sensitivity, establishing DSI as a practical tool for precise and safe cryoablation monitoring.

## Acknowledgments

This work was supported in part by the Israel Science Foundation under Grant 192/22, in part by an ERC StG under Grant 101041118 (NanoBubbleBrain), in part by the Israel Cancer Research Fund (grant number 1286686), and in part by the Nicholas and Elizabeth Slezak Super Center for Cardiac Research and Biomedical Engineering at Tel Aviv University.

## Contributions

G.L. designed and performed the research, conducted experiments, analyzed the data, and wrote the manuscript. T.G. guided and advised the analysis process. M.B. assisted with system setup and data acquisition. T.I. guided, advised, and designed the research and wrote the manuscript. All authors reviewed the manuscript.

